# Translational regulation of *Syngap1* by FMRP modulates NMDAR mediated signalling

**DOI:** 10.1101/345058

**Authors:** Abhik Paul, Bharti Nawalpuri, Shruthi Sateesh, Ravi S Muddashetty, James P Clement

## Abstract

SYNGAP1, a Synaptic Ras-GTPase activating protein, regulates synapse maturation during a critical developmental window. Heterozygous mutation in *SYNGAP1* (*SYNGAP1*^+/-^) has been shown to cause Intellectual Disability (ID) in children. Recent studies have provided evidence for altered neuronal protein synthesis in a mouse model of *Syngap1*^+/-^. However, the molecular mechanisms behind the same is unclear. Here, we report the reduced expression of a known translation regulator, FMRP, during a specific developmental period in *Syngap1*^+/-^ mice. Our results demonstrated that FMRP interacts with and regulates the translation of *Syngap1* mRNA. We further show that, during development, reduced translation of FMRP and this decrease in FMRP leads to a compensatory increase of *Syngap1* translation in *Syngap1*^+/-^. These developmental changes are reflected in the altered response of eEF2 phosphorylation downstream of NMDA receptor signalling. We propose a cross-talk between FMRP and SYNGAP1 mediated signalling which can also explain the compensatory effect of impaired signalling observed in *Syngap1*^+/-^ mice.

## Introduction

SYNGAP1 is a synaptic RAS-GTPase Activating Protein (SYNGAP1), which acts downstream of N-Methyl D-Aspartate Receptors (NMDAR), and negatively regulates RAS GTPase (Kim, Liao et al. 1998, Komiyama, Watabe et al. 2002). NMDAR activation leads to phosphorylation of SYNGAP1 by Ca^2+^/Calmodulin-dependent Kinase II (CAMKII) (Krapivinsky, Medina et al. 2004). Phosphorylated SYNGAP1 is rapidly dispersed from dendritic spine to dendritic shaft leading to activation of its downstream molecules in dendritic spines (Araki, Zeng et al. 2015). Removal of SYNGAP1 from spine leads to increased activity of Extracellular Signal Regulated Kinases (ERK) via RAS (Rumbaugh, Adams et al. 2006), which further allows insertion of α-amino 3-hydroxy 5-methyl 4-isoxazolepropionic acid Receptors (AMPAR) on the post-synaptic membrane (Zhu, Qin et al. 2002).

Using mouse model, studies have shown that *Syngap1*^+/-^ causes early maturation of dendritic spines in the hippocampus (Clement, Aceti et al. 2012) and altered critical period of development in Barrel cortex (Clement, Ozkan et al. 2013). Furthermore, these studies have shown altered excitatory synapaptic activity and AMPA/NMDA occurs during Post-Natal Day (PND)14-16 in the hippocampus of *Syngap1*^+/-^ mice. Consistent with its molecular function, studies from human patients have shown that loss of function mutations in *SYNGAP1* results in Intellectual Disability (ID) and epileptic phenotype (Hamdan, Gauthier et al. 2009, Hamdan, Daoud et al. 2011, Rauch, Wieczorek et al. 2012). All these studies suggest that SYNGAP1 is crucial for the development of excitatory circuit during a critical period of development (Jeyabalan and Clement 2016)

Recent studies using *Syngap1*^+/-^ mice and *Syngap1* knock-down in rat cultured cortical neurons demonstrated increased levels of basal protein synthesis in *Syngap1*^+/-^ as compared to Wild-type (Wang, Held et al. 2013, Barnes, Wijetunge et al. 2015). The study also suggested that SYNGAP1 modulates insertion of AMPARs at the post-synaptic membrane, thereby regulating synaptic plasticity through protein synthesis (Rumbaugh, Adams et al. 2006, Wang, Held et al. 2013). However, the molecular mechanisms of SYNGAP1-medaited regulation of protein synthesis, particularly during development, is unclear.

To regulate synaptic protein synthenis, SYNGAP1 may crosstalk with other translation regulators. One such potential candidate to consider is FMRP. Similar to *Syngap1*^+/-^ mice, *Fmr1* KO results in exaggerated levels of basal protein synthesis and altered dendritic spine structure and function (Huber, Gallagher et al. 2002). Additionally, a recent report showed exaggerated protein synthesis-independent mGluR-LTD (Metabotropic glutamate receptor-dependent long-term depression) in *Syngap1*^+/-^ (Barnes, Wijetunge et al. 2015), which is another hallmark phenotype of FMRP associated synaptic deficit (Huber, Gallagher et al. 2002). Based on these findings, we hypothesise a possible cross-talk between SYNGAP1 and FMRP in regulating activity mediated protein synthesis at the synapse. In this study, we have shown that FMRP level was altered during development, especially in PND21-23, in *Syngap1*^+/-^. In addition, FMRP interacts with and regulates the translation of *Syngap1* mRNA, and, thus, compensates for *Syngap1* translation in *Syngap1*+/-. These results may explain the impaired NMDAR-mediated signalling observed in *Syngap1*^+/-^.

## Materials and Methods

### Animals

C57/BL6 WT and *Syngap1^+/-^* mice were obtained from The Jacksons Laboratory (https://www.jax.org/strain/008890) (Kim, Lee et al. 2003) and bred and maintained in the Institute’s animal house under 12-hour dark and light cycle. This study was carried out in accordance with the principles of the Basel Declaration and recommendations of the Institutional Animal Ethics Committee (IAEC; Prof Anuranjan Anand, Chairman). The protocol was approved by the Committee for Purpose of Control and Supervision of Experiments on Animals (CPCSEA; Dr. K.T. Sampath, CPCSEA Nominee)

### Preparation of hippocampal slices

Acute brain slices were prepared from PND>90 male and female Wild type (WT) and *Syngap1^-/+^* mice. All mice were bred and maintained in the Animal facility, JNCASR. Mice were brought from the animal house and sacrificed by cervical dislocation, and brain was dissected out. The brain was kept in ice cold sucrose based artificial cerebrospinal fluid (aCSF; cutting solution) comprising: 189 mM Sucrose (S9378, Sigma), 10 mM D-Glucose (G8270, Sigma), 26 mM NaHCO_3_ (5761, Sigma), 3 mM KCl (P5405, Sigma), 10 mM MgSO_4_.7H_2_O (M2773, Sigma), 1.25 mM NaH_2_PO_4_ (8282, Sigma) and 0.1 mM CaCl_2_ (21115, Sigma). Brain was taken out of cutting solution and glued to the brain holder of the vibratome (Leica #VT1200), and 350 μm thick horizontal slices were prepared. Cortex was dissected out from each slice to obtain only the hippocampus. All the slices were kept in slice chamber containing aCSF comprising: 124 mM NaCl (6191, Sigma), 3 mM KCl (P5405, Sigma), 1 mM MgSO_4_.7H_2_O (M2773, Sigma), 1.25 mM NaH_2_PO_4_ (8282, Sigma), 10 mM D-Glucose (G8270, Sigma), 24 mM NaHCO_3_ (5761, Sigma), and 2 mM CaCl_2_ (21115, Sigma) in water bath (2842, ThermoScientific) at 37°C for 40-45 minutes. Following recovery, slices were kept in room temperature (RT) till the experiment completed. Post dissection, every step was carried out in the presence of constant bubbling with carbogen (2-5% CO_2_ and 95% O_2_; Chemix, India). All measurements were performed by an experimenter blind to the experimental conditions.

### Extracellular Field Recordings

Field excitatory post synaptic potential (fEPSP) were elicited from pyramidal cells of CA1 regions of stratum radiatum by placing concentric bipolar stimulating electrode (CBARC75, FHC, USA) connected to a constant current isolator stimulator unit (Digitimer, UK) at Schaffer-collateral commissural pathway, and recorded from stratum radiatum of CA1 area of the hippocampus, with 3-5 MΩ resistance glass pipette (ID: 0.69 mm, OD: 1.2 mm, Harvard Apparatus) filled with aCSF. Signals were amplified using Axon Multiclamp 700B amplifier (Molecular Devices), digitized using an Axon Digidata 1440A (Molecular Devices), and stored on a personal computer. Online recordings and analysis were performed using pClamp10.7 software (Molecular Devices). Stimulation frequency was set at 0.05 Hz. mGluR-LTD was induced by 5 minutes bath application of the Group I mGluR agonist (*S*)-3,5-dihydroxyphenylglycine (DHPG; Cat# 0805, Tocris, UK).

### Lysate Preparation

Brain lysates were prepared from Post-Natal Day (PND) 4-5, 7-9,14-16, 21-23, and adults (2-5 months). WT and *Syngap1^+/-^* mice were sacrificed by cervical dislocation, brain was dissected out, and hippocampus was separated in ice cold Phosphate Buffered Saline (PBS) of pH 7.4 containing NaCl (137 mM, S6191, Sigma), KCl (2.7 mM, P5405, Sigma), Na_2_HPO_4_ (10 mM, 10028-24-7, Fisher Scientific), KH_2_PO_4_ (1.8 mM, GRM1188, HIMEDIA). The tissue was homogenised using RIPA buffer containing NaCl (150 mM, S6191, Sigma,), Tris-HCl (50 mM, Tris: 15965, Fisher Scientific; HCl: HC301585, Merck) pH 7.4, EDTA (5 mM, 6381-92-6, Fisher Scientific), Na-Deoxycholate (0.25%, Triton X (RM 845, HiMedia). Additionally, 1X Protease Inhibitor (P5726, Sigma,), and 1X Phosphatase Inhibitor Cocktail 2 and 3 (P5726 and P0044 respectively, Sigma) was added to the buffer to increase the stability of the lysate. Then, the homogenates were centrifuged at 16000g for 30 minutes at 4°C. The samples were aliquoted and stored at −80°C. The supernatants were collected, and the protein was estimated using Bradford (5000006, Biorad) assay.

### SDS-PAGE and Western Blotting

The protein samples were electrophoresed onto SDS (161-0302, Biorad) Polyacrylamide (161-0156, Biorad) 5% stacking gel for 30 minutes and 8% resolving gel (for FMRP, SYNGAP1, MOV10, PSD95) for around 2 hours or 10% resolving gel (Phospho-eEF2, Total-eEF2 and RPLP0) for around 3 hours. The proteins were transferred for 3 hours at 80V at 4°C onto Polyvinylidene Fluoride (PVDF) membrane (1620177, Bio-Rad) and blocked using 5% skimmed milk (GRM 1254, HIMEDIA) or 5% BSA (GRM105, HIMEDIA) in PBS for 1-hour in Room Temperature (RT) at 25°C. BSA was used for blocking of all Phospho-Proteins. For MOV10, blocking was done for ~3-4 hours at RT. The blots were washed with 1% PBST (PBS+ Tween 20; GRM156 HIMEDIA) for 3 times for 10 minutes, and incubated with Primary Antibodies for FMRP (F4055, Sigma Aldriech, 1:1000 dilution, raised in rabbit), β-Actin (PA116889, Thermofischer, 1:15000 dilution, raised in rabbit), and MOV10 (ab80613, Abcam, 1:1000 dilution, raised in rabbit), PSD95 (MA1-046, Thermofisher, 1:1000 dilution, raised in mouse), Phospho-eEF2 (Thr 56) 2331S, Cell Signalling Technology, 1:1000 dilution, raised in rabbit), Total-eEF2 (2332S, Cell Signalling Technology, 1:1000 dilution, raised in rabbit), and RPLP0 (ab101279, abcam, 1:1000 dilution, raised in rabbit) overnight. After primary incubation, blots were given PBST wash for 3 times for 10 minutes, then incubated with anti-Rabbit (1706515, Biorad) / anti-Mouse (1706516, Biorad) HRP conjugated Secondary antibody (1:10000 dilution). After subsequent washes with PBST, the blots were developed by chemiluminescent method using ECL western clarity solution (1705060, BioRad). Image was taken in Versa Doc (Biorad) or in ImageQuant (LAS 4000 from GE) further merged using ImageLab version 5.2.1 and bands were quantified using ImageJ software.

### Cell culture and transfection

HeLa cells were maintained in DMEM containing 10% FBS at 37°C in a 5% CO_2_ environment passaged using 0.05% trypsin-EDTA solution. Transfections were performed using lipofectamine 2000 (11668027, Thermofisher) as per manufacturer’s protocol.

### Polyribosome Profiling assay

Hippocampus were dissected out from Post-natal day (PND)21-23 and PND14-16 *Syngap1^+/-^* (HET) and Wild-type (WT; littermates) as described earlier. Tissue was homogenised using Lysis buffer containing Tris-HCl (200 mM, Tris: 15965, Fisher Scientific; HCl: HC301585, Merck), KCl (100 mM, P5405, Sigma), MgCl_2_ (5 mM, M8266, Sigma), Dithiothreitol (DTT, 1 mM, 3483-12-3, Sigma), NP40 (1%), and 1X Protease Inhibitor cocktail (P5726, Sigma). All reagents were dissolved in Diethylpyrocarbonate (DEPC, D5758, Sigma) treated autoclaved water. Samples were aliquoted into two equal parts. One part was treated with Cycloheximide (CHX, 10 μg/ml, C7698, Sigma), and another part was treated with Puromycin (1 mM, P9620, Sigma), and both are translation inhibitors. After CHX and Puromycin addition, the lysates were kept at 37°C for 30 minutes and centrifuged at 4°C for 30 minutes at 18213g. Supernatant was further loaded carefully on the sucrose gradient prepared in polysome tubes. Sucrose (84097, SIGMA) gradient tubes were prepared 1-day prior to the day of the experiment. 15% to 60% gradients were made, and stored at −80°C. The samples (supernatants) were added to each polysome tubes (331372, BECKMAN COULTER) slowly, and ultra-centrifuged (Beckman, OptimaXL 100K) at 4°C at 39000 RPM for 1 hour and 40 minutes. The tubes were then transferred to UV Visible spectrophotometer (Model: Type 11 Optical unit with reference Flowcell/No bracket, Serial No: 213K20162 at National Centre for Biological Sciences (NCBS)), and Fraction collector instrument (from TELEDYNE ISCO at NCBS). Fractions were collected at A_254_ spectra. The bottom of the tube was pierced using needle syringe attached to a pipe holding 60% sucrose. Addition of 60% sucrose from the bottom created a positive pressure, which pushed the fractions to come out from the top of the polysome tube. Then, the fractions were collected in 1.5 ml tubes. Total of 11 fractions were collected from each tube and these fractions were treated with SDS loading dye containing β-Mercaptoethanol (MB041, HIMEDIA). SDS-PAGE was done for these fractions and immunoblotted for RPLP0 protein.

### Synaptoneurosome preparation and NMDA stimulation

The Hippocampus were dissected out as described earlier. Mouse hippocampus was homogenised in 1000 μl of Synaptoneurosome buffer containing NaCl (116.5 mM, S6191, Sigma), KCl (5 mM, P5405, Sigma), MgSO_4_ (1.2 mM, M7506, Sigma), CaCl_2_ (2.5 mM, C5670, Sigma), KH_2_PO_4_ (1.53 mM, GRM1188, HIMEDIA), Glucose (3.83%, G8270, Sigma), 1X Protease Inhibitor Cocktail (P5726, Sigma). Homogenate was filtered through 3, 100 μm filter (NY1H02500, Merck Millipore), then through 11 μm filter once (NY1102500, Merck Millipore). The filtrate obtained was centrifuged at 1500g for 15 minutes at 4°C. Pellet was resuspended in 1 ml Synaptoneurosome buffer. NMDA receptor stimulation was done by applying NMDA (40 μM, M3262, Sigma) for 1- and 2-minutes respectively at 37°C in 350 RPM. After stimulation, the synaptoneurosome was centrifuged at 11000 RPM for 21 seconds, and pellet was resuspended in Lysis buffer followed by centrifugation at 18213g at 4°C for 30 minutes. The supernatant was taken and denatured in SDS and β-Mercaptoethanol (MB041, HIMEDIA), and western blot was done.

### RNA extraction and qPCR

Total RNA was extracted from the polysome fractions by Trisol (15596026, Thermofisher Scientific) method and the mRNAs were converted to cDNA using iScript cDNA synthesis kit (1708891, Biorad). qPCR was performed for *Syngap1*, *Fmr1*, and *β-actin* (Primers were designed and obtained from Sigma) using CFX384 Real Time System from Biorad. SYBR green was obtained from Biorad (1725122). Ct values obtained from the reactions were converted to the copy number of the mRNA, and then percentage of these copy numbers in each fraction were plotted. List of primers used are mentioned below.

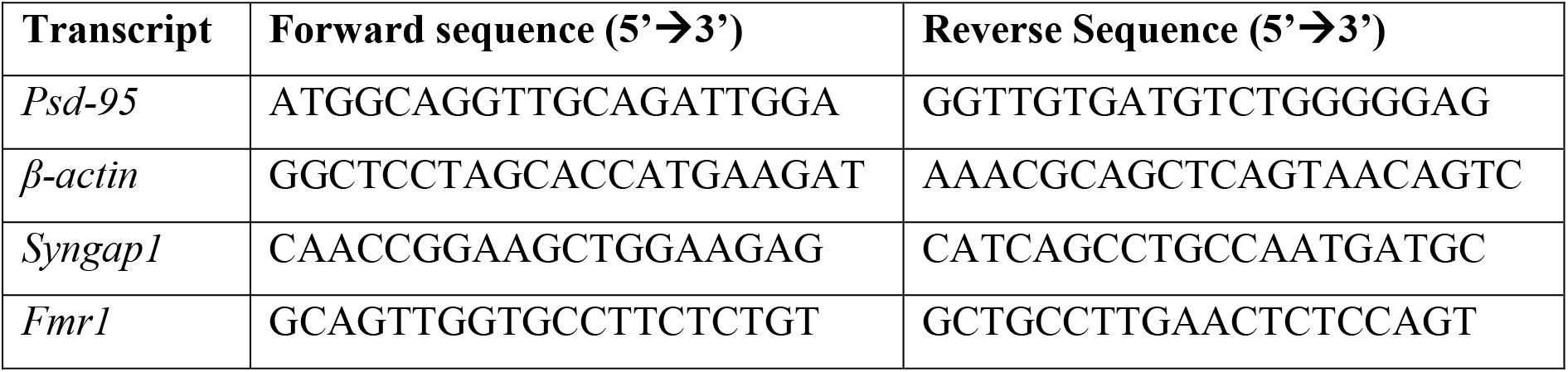

### Statistics

All graphs were plotted in Graph Pad Prism 7, and Microsoft Excel 2016. Extracellular field recordings were performed and analysed using Clampfit10.7, and Excel 2016. Time course data were plotted by averaging every 2 minutes. Example traces are those recorded for 1-2 min around the time point indicted. Error bars correspond to ±SEM (Standard Error of Mean). Unpaired Student’s *t-*test and 2-way ANOVA were performed to test for difference between groups and different age, unless otherwise stated.

## Results

### Reduced FMRP level during development in *Syngap1*^+/-^

Studies have shown that Group I mGluR and NMDA receptors interact via Homer-Shank that may regulate protein synthesis (Tu, Xiao et al. 1999, Bertaso, Roussignol et al. 2010). To determine whether Group I mGluR activation in *Syngap1^-/+^* results in altered protein synthesis and hippocampal pathophysiology similar to *Fmr1^-/y^*, Group I mGluR-mediated LTD (mGluR-LTD) was induced in the Schaffer-collateral pathway in adult mice by bath applying 50 μm (*S*)-DHPG, Group I mGluR agonist, for 5 minutes. We observed significantly increased mGluR-LTD in *Syngap1^+/-^* mice (*Syngap1*^+/-^ referred as HET in Figures; 47±4% LTD) as compared to their Wild-type littermate controls (61±3% LTD; p=0.012; **Fig 1A**). This result suggests that mGluR-LTD in *Syngap1*^+/-^ is similar to *Fmr1^-/y^* as shown by Barnes et al. (Barnes et al., 2015), and further reiterates the fact that protein synthesis downstream of synaptic signalling is likely to be altered in *Syngap1*^+/-^ similar to *Fmr1^-/y^*, which prompted us to investigate the role of FMRP in the pathophysiology of *Syngap1^+/-^* mutation.

**Figure 1:**
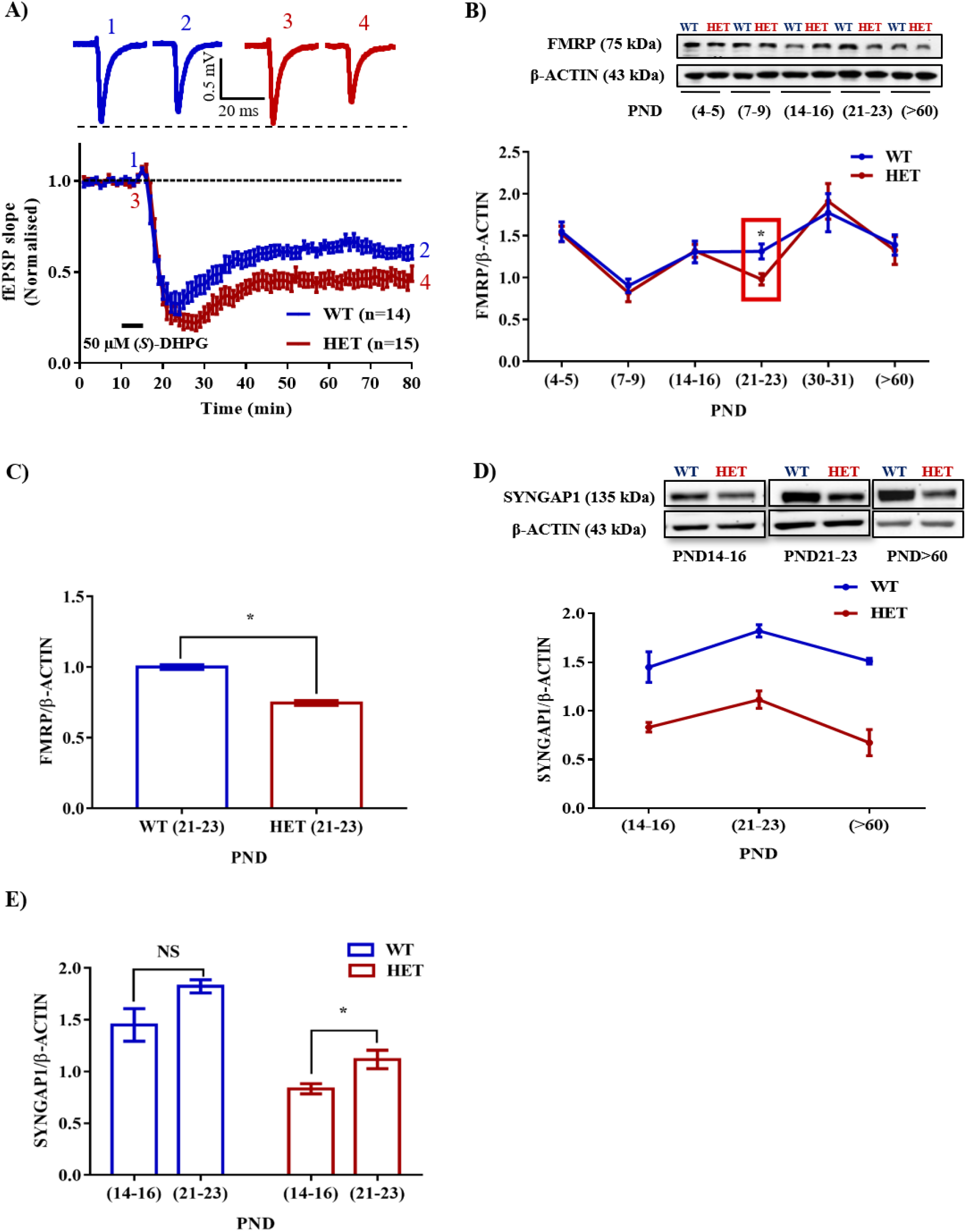
Altered expression of FMRP in the Hippocampus of *Syngap1*^+/-^ during development. **A.** Application of 50 μm (*S*)-DHPG induced enhanced Group I mGluR mediated LTD in the Schaffer-collateral pathway of *Syngap1*^+/-^ (HET) compared to Wild-type (WT) littermates. Sample traces obtained before and after induction of LTD as indicated by time points (*top*). WT=61±3% LTD, n=14; HET=47±4% LTD, n=15; * p<0.05 **B.** Representative immunoblot for FMRP level in the hippocampus during development (*top*). Pooled data of FMRP level normalised to β-ACTIN in the hippocampus during development (*below*). PND4-5 (WT: n=12; HET: n=7), PND7-9 (WT: n=8; HET: n=7), PND14-16 (WT: n=10; HET: n=8), PND21-23 (WT: n=14; HET: n=10), PND30-31 (WT: n=4; HET: n=7), PND>60 (WT: n=8; HET: n=10). *p<0.05; Two-way ANOVA, F _(*5, 93*)_ = 0.7965. **C.** Bar graph derived from **1B** depicting FMRP level during PND21-23 in WT (n=14) and HET (n=10), *p<0.05, NS = not significant. **D.** Representative Immunoblots for SYNGAP1 during development (*top*). SYNGAP1 level normalised to β-ACTIN during PND14-16 (WT: n=5; HET: n=5), PND21-23 (WT: n=4; HET: n=5), and adults *i.e*, PND>60 (WT: n=4; HET: n=3); ****P<0.0001, Two-way ANOVA, F _(*1, 15*)_ = 0.1893. **E.** Bar graph derived from **1D** shows increased SYNGAP1 level in HET during PND21-23 (WT: n=4; HET: n=5) when compared to PND14-16 (WT: n=5; HET: n=5) while no significant change was observed in WT. *p<0.05, NS= not significant.

We studied the expression of FMRP in the hippocampus of Wild-type and *Syngap1^+/-^* mice during different post-natal days, starting from PND4 till 2-5 months old. Using quantitative immunoblotting, we observed that FMRP level (normalised to β-ACTIN) was reduced in *Syngap1^+/-^* mice (0.98±0.06) as compared to Wild-type in PND21-23 (1.31±0.09; p=0.0120; **Fig 1B-C**) but not in other age groups. Previous studies have shown that SYNGAP1 level was altered during development in *Syngap1^+/-^* mice that affected synaptic transmission (Vazquez, Chen et al. 2004, Clement, Aceti et al. 2012). To study whether reduced level of FMRP is compensating for the altered SYNGAP1 level in *Syngap1^+/-^* mice, expression of SYNGAP1 in Wild-type and *Syngap1^+/-^* mice was quantified as shown in **Fig 1D (**Genotype: p<0.0001). Upon further analysis, we found that SYNGAP1 level was increased during PND21-23 (1.12±0.09) compared to PND14-16 in *Syngap1^+/-^* (0.83±0.05; p=0.0236; **Fig 1E**). In contrast, the level of SYNGAP1 was not altered significantly between PND21-23 (1.82±0.06) and PND14-16 (1.33±0.08) in Wild-type mice (p=0.0863; **Fig 1E**).

### FMRP interacts with *Syngap1* mRNA and regulates its translation

FMRP is a known regulator of synaptic translation (Osterweil, Krueger et al. 2010). A previous study using HITS-CLIP has reported that one of the FMRP targets is *Syngap1* mRNA (Darnell & Klann, 2013). G-quadruplexes are one of the structures present in RNA which can be recognised by FMRP (Darnell, Jensen et al. 2001). Bioinformatics analysis using Quadraplex forming G-Rich Sequences (QGRS) Mapper predicted the presence of multiple G-quadruplexes structures with high G-Score in *Syngap1* mRNA (**Supplementary Fig 1A**). In addition, G-quadruplex forming residues were found to be conserved among mice, rat, and human *Syngap1* mRNA (**Supplementary Fig 1B**). To further confirm the interaction of FMRP with *Syngap1* mRNA, we performed FMRP immunoprecipitation from mouse hippocampal lysates to investigate the enrichment of *Syngap1* mRNA by qPCR. We observed ~2-fold enrichment of *Syngap1* mRNA relative to *β-actin* mRNA in FMRP IP pellet over supernatant (2.16±0.18, p=0.0036; **Fig 2A**). *Psd-95* mRNA, a known FMRP target mRNA (Muddashetty, Nalavadi et al. 2011) showed a significant 3.5-fold enrichment compared to *β-actin* mRNA (p=0.0001; 3.59±0.07; **Fig 2A**) which we used as a positive control. These results demonstrate that FMRP interacts with *Syngap1* mRNA.

**Figure 2.**
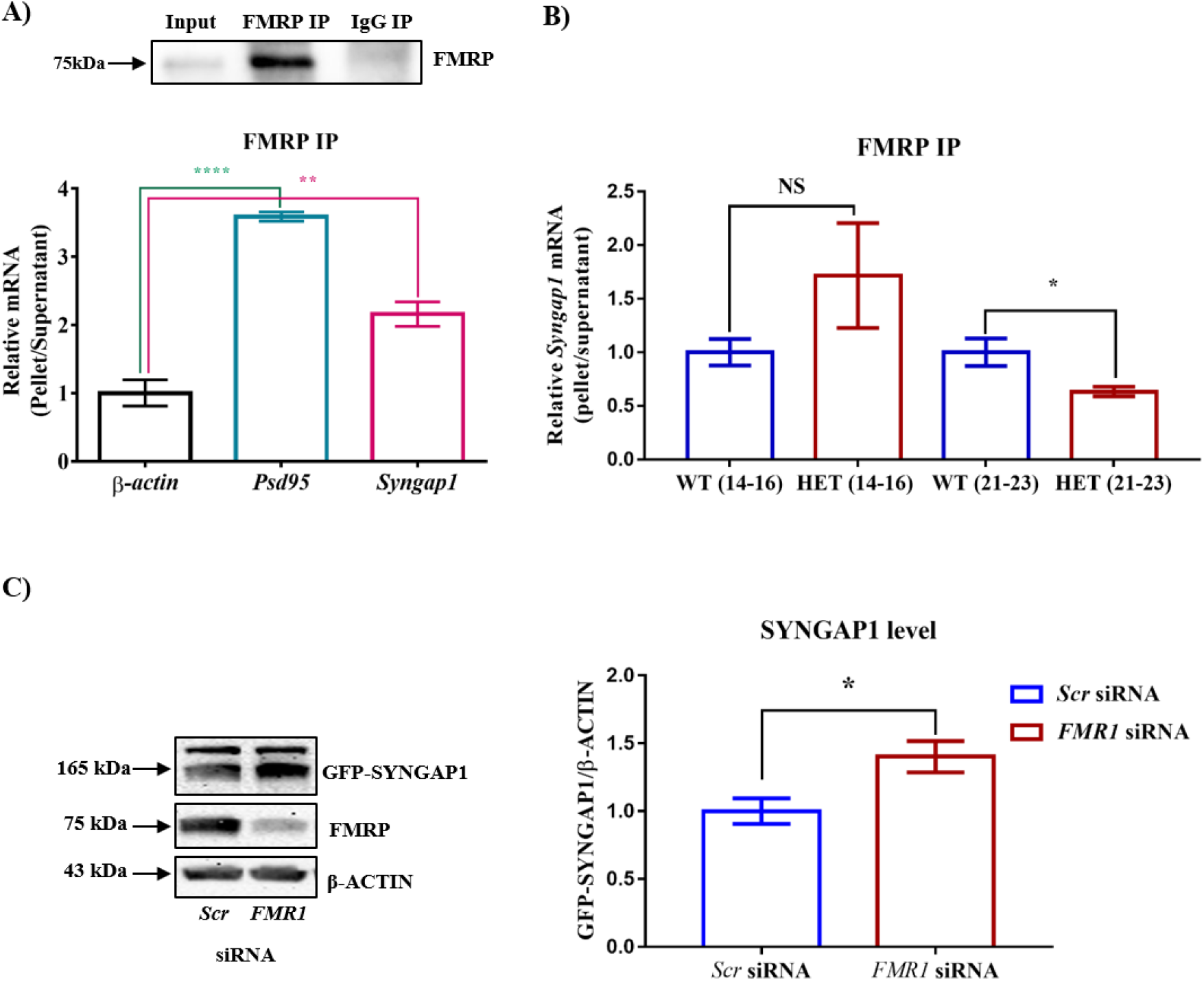
FMRP regulates *Syngap1* mRNA translation. **A.** Immunoblot for FMRP following FMRP-IP and IgG-IP (*top)*. Bar graph showing relative *Syngap1*, *Psd95* mRNA enrichment in FMRP IP pellet compared to Supernatant after normalising to *β-Actin* (WT: n=3; Below). One-way ANOVA followed by Dunnett’s multiple comparisons test. **p<0.01, ****p<0.0001, **B.** Bar graph shows relative *Syngap1* mRNA enrichment in FMRP IP pellet compared to supernatant from hippocampus at PND21-23 (WT: N=5; HET: N=4) and PND14-16 (WT: N=7; HET: N=3) normalised to WT. Unpaired Student’s *t*-test; *p<0.05; NS= not significant. **C.** Representative immunoblot for SYNGAP1, FMRP and β-ACTIN showing knock-down of FMRP leads to increase SYNGAP1 expression in Hela (*left*). Quantified bar graph shows increase in the level of GFP-SYNGAP1 expression in the cells treated with *FMR1* siRNA compared to *Scr* SiRNA treatment (*right*). Unpaired Student’s *t*-test; *p<0.05.

Interaction between FMRP and *Syngap1* mRNA was decreased in *Syngap1^+/-^* (p=0.045; 0.63±0.04; **Fig 2B**) compared to WT (1.0±0.13) at PND21-23. Whereas in PD14-16, we observed a trend of increased interaction though we did not find statistical significance (p=0.28; *Syngap1^+/-^*=2.3±0.66; WT=1.0±0.1; **Fig 2B**). We did not observe any change in the interaction of *Psd-95* mRNA with FMRP at any of these age groups (PND14-16: p=0.44; *Syngap1^+/-^* =1.297±0.34; WT=1.0±0.2; PND21-23: p=0.24; *Syngap1^+/-^*=0.8347±0.12; WT=1.0±0.07; **Supplementary Fig 2A**) Further, we overexpressed *GFP*-*Syngap1* in Hela cells followed by knock down of *FMR1* (decreased expression of FMRP; **Supplementary Fig 2B, 2C, 2D**). We show that reduction in FMRP led to increased level of GFP-SYNGAP1 (p=0.01; *Scr* SiRNA 0.58±0.05; *FMR1* SiRNA 0.82±0.067; **Fig 2C**). These results show FMRP not only interacts with *Syngap1* mRNA but also regulates its translation. In addition, our data demonstrates that reduced interaction in *Syngap1^+/-^* at PND21-23 might lead to increased SYNGAP1 level.

### *Syngap1* mRNA translation is differentially regulated in *Syngap1^+/-^*

To further understand the compensatory increase in SYNGAP1 levels during PND21-23 in *Syngap1^+/-^*, we analysed *Syngap1* mRNA translation status at PND14-16 and PND21-13. We studied translation by Polysome profile (**Fig 3A**), from hippocampal lysates of Wild-type and *Syngap1^+/-^* mice at PND14-16 and PND21-23 (Muddashetty, Kelic et al. 2007). **Fig 3B** demonstrates that the A_254_ traces from cycloheximide-treated samples curve showed the distinct peaks corresponds to mRNP, monosome, and polysomes respectively. A_254_ traces between Wild-type and *Syngap1^+/-^* mice did not show any significant difference, suggesting that the global translation in hippocampus might be unaffected in *Syngap1^+/-^* mice. Further, immunoblots for Ribosomal large subunit protein, RPLP0, has shown a shift in puromycin treated sample as puromycin disassemble the ribosome from translating mRNA (**Fig 3B**), along with a shift in *β-actin* mRNA (**Supplementary Fig 3A**). In our experiments, fraction numbers 1 to 6 and 7 to 11 were considered as non-translating fractions and translating fractions or polysome (puromycin-sensitive) respectively (**Supplementary Fig 3A,3B**). Further, we estimated the RPLP0 distribution in translating/non-translating fractions of WT and *Syngap1*^+/-^ during PND14-16 (WT=1.06±0.18, *Syngap1*^+/-^=0.71±0.15, p=0.22) and PND21-23 (WT=1.27±0.21, *Syngap1*^+/-^=0.83±0.17, p=0.14), suggesting no significant change in the distribution of RPLP0 (**Fig 3C, Supplementary Fig 3C**). To understand the translation status of *Syngap1* mRNA during PND14-16 and PND21-23, we quantified *Syngap1* mRNA present in translating fraction by performing quantitative PCR from RNA isolated from both non-translating (Fraction 1-6) and translating fraction (Fraction 7-11). However, a significant reduction in translating *Syngap1* mRNA in *Syngap1^+/-^* mice during PND14-16 compared to WT (WT=84.32±4%; *Syngap1^+/-^*=65.77±2%; p=0.0018; **Fig 3D**) was observed. On the contrary, this difference was absent during PND21-23 (WT=92.9±3.5%; *Syngap1^+/-^* =87.9±2.5%; p=0.3033). As a control, distribution of *β-actin* mRNA in translating pool was quantified, and no significant difference was observed between Wild-type and *Syngap1^+/-^* at PND14-16 (WT=89.9±3%; *Syngap1^+/-^*=83.6±2.9%; p=0.2039) and PND21-23 (WT=97.9±0.6%; *Syngap1^+/-^*=89.8±3.8%; p=0.0697; **Supplementary Fig 3B**). These results suggest that an increase in *Syngap1* mRNA translation leads to the corresponding increase in SYNGAP1 level observed in PND21-23 when compared to PND14-16 in *Syngap1^+/-^*.

**Figure 3.**
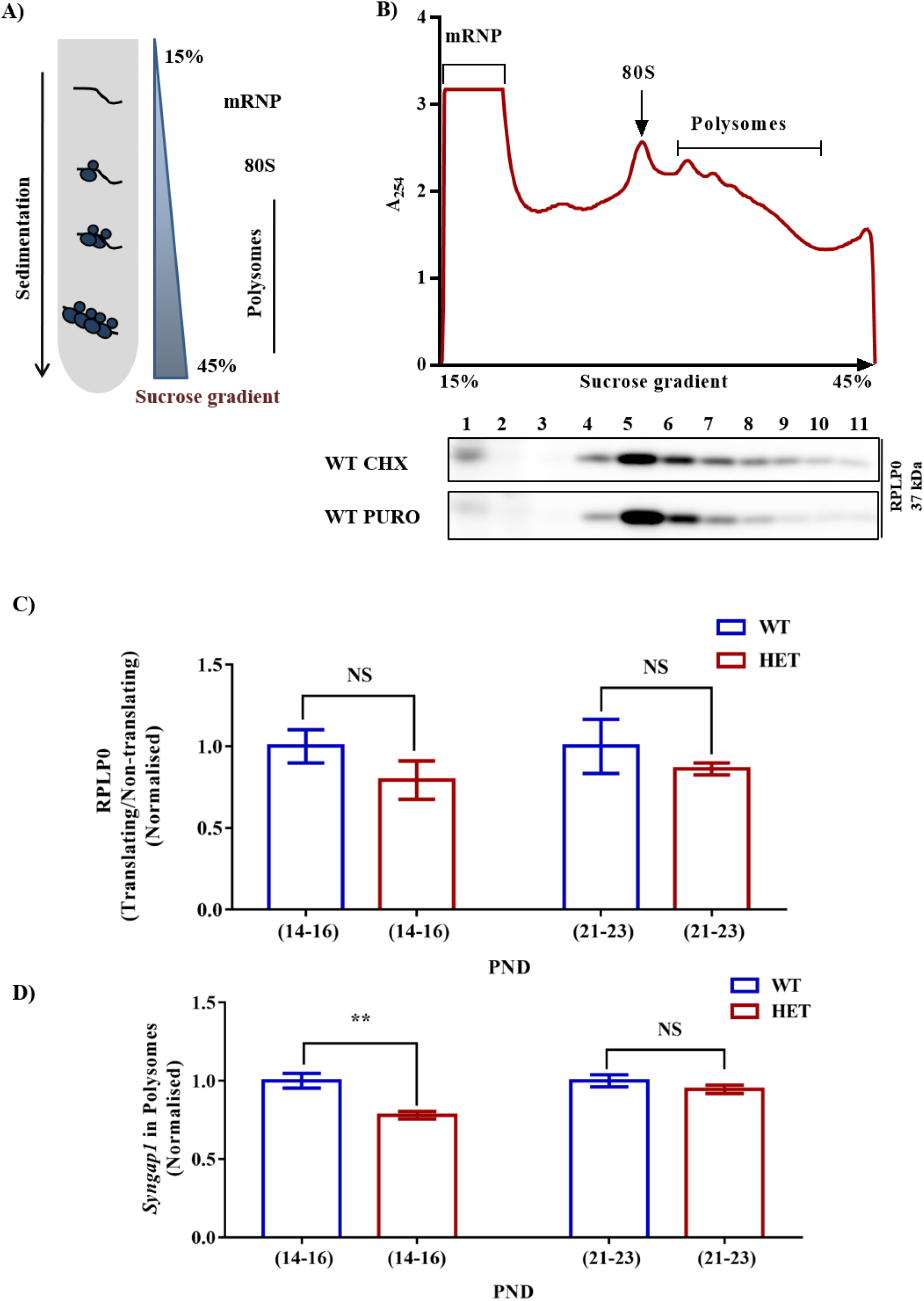
Altered *Syngap1* mRNA distribution in translating polysomal fractions of HET. **A.** Schematic diagram depicting sucrose gradient method used for polyribosome profiling (translation assay) **B.** Polyribosome profile obtained from Cycloheximide treated hippocampal lysate during PND14-16 in HET (*top*). Representative immunoblots for RPLP0 distribution in Cycloheximide and Puromycin treated polysome during PND14-16 (*below*). **C.** Bar graph shows RPLP0 distribution in Translating/Non-translating fractions during PND14-16 (WT: n=5; HET: n=3; p=0.22) and PND21-23 (WT: n=4; HET: n=4; p=0.14). Unpaired Student’s *t*-test was done for both age groups. NS = not significant. **D.** *Syngap1* mRNA distribution in polysome in HET normalised to WT during PND14-16 (WT: n=4; HET: n=6; p=0.0018) and PND21-23 (WT: n=3; HET: n=3; p=0.3033).

### Reduced FMRP in polysome at PND21-23 in *Syngap1*^*+/-*^

To understand if the changes in the levels of translating *Syngap1* mRNA is a result of the altered association of FMRP with polysomes, we estimated the distribution of FMRP in translating/non-translating fraction from polysome profiling. We observed that the level of FMRP was increased in PND14-16 in the polysomal fraction in *Syngap1^+/-^* (0.41±0.03) as compared to age-matched Wild-type (0.17±0.02) control (p=0.0011; **Fig 4A, 4B**). However, we observed reduced FMRP distribution in the polysomal fraction of *Syngap1^+/-^* (0.23±0.03) in PND21-23 compared to Wild-type (0.55±0.15; p=0.0473; **Fig 4A, 4B**). This might have a compounding effect on translation of FMRP target mRNAs during PND21-23 in *Syngap1^+/-^* as the overall FMRP level was also reduced. As a control, we analysed the PSD-95 levels during PND14-16 (WT=2.03±0.35; *Syngap1^+/-^*=1.39±0.15; p=0.125) and PND21-23 (WT=0.98±0.05; *Syngap1^+/-^*=0.98±0.12; p=0.96). However, we did not observe any change between Wild-type and *Syngap1^+/-^* (**Supplementary Fig 4A, 4B**).

**Figure 4.**
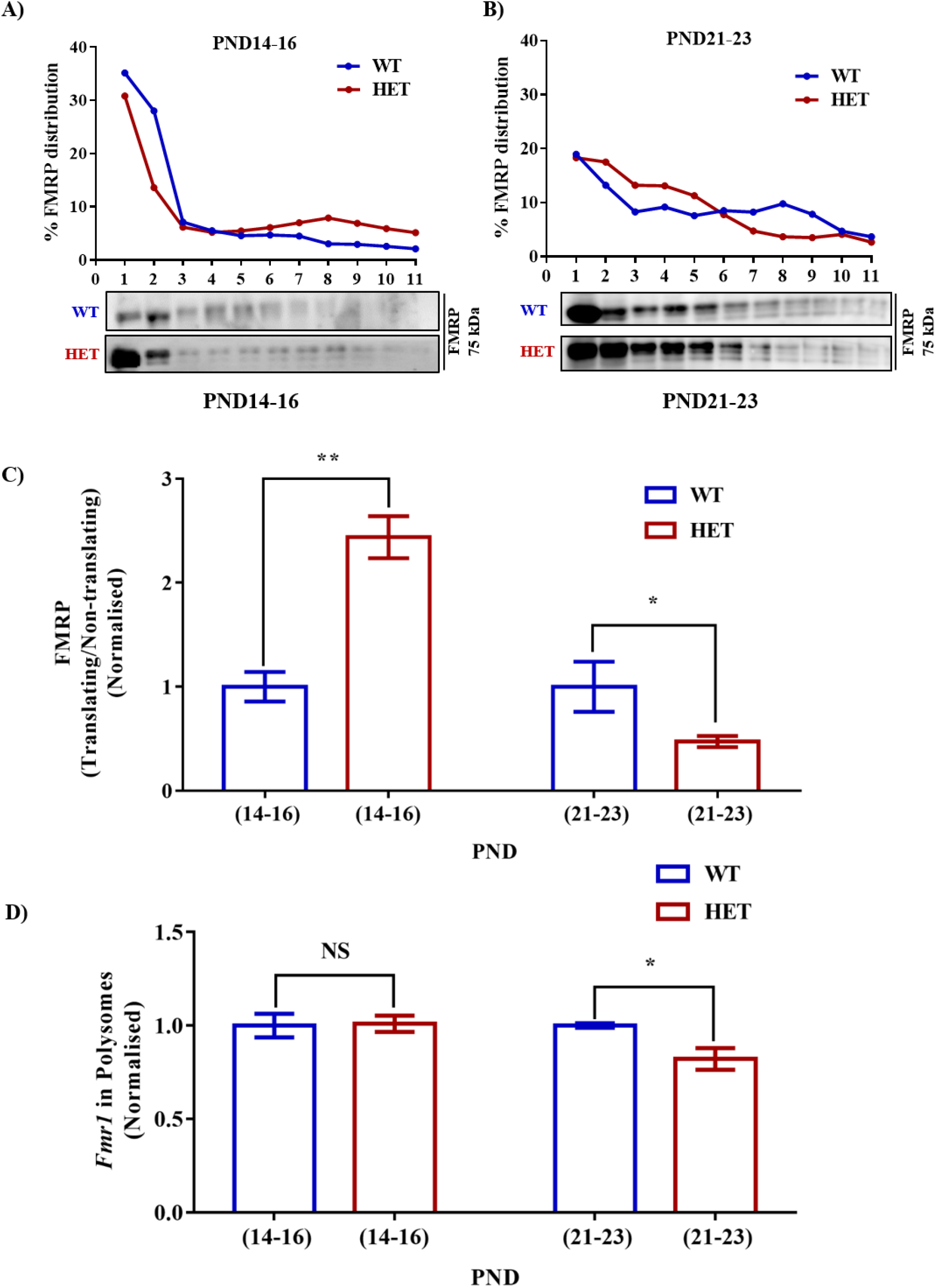
Decreased *Fmr1* mRNA and FMRP distribution in translating fractions of polysomes in HET during PND21-23. **A.** Representative line graphs showing percentage FMRP distribution in polysomes during PND14-16 (*top*) along with representative immunoblot for FMRP distribution (*below*). **B.** Line graph showing representative percentage FMRP distribution in polysomes during PND21-23 (*top*) and the corresponding representative immunoblot for FMRP distribution (*below*). **C.** Bar graph showing FMRP distribution in translating/non-translating fractions in HET normalised to WT during PND14-16 (WT: n=4; HET: n=4) and PND21-23 (WT: n=4; HET: n=5). *p<0.05, **p<0.01. **D.** Bar graph depicting relative *Fmr1* mRNA in translating fractions of HET normalised to WT during PND14-16 (WT: n=3; HET: n=3) and PND21-23 (WT: n=3; HET: n=5). *p<0.05, NS= not significant.

Our previous result showed reduced FMRP level during PND21-23 in *Syngap1^+/-^* as compared to its Wild-type counterpart (**Fig 1C**). We investigated whether the reduced level of FMRP is due to altered *Fmr1* mRNA levels or translation. We evaluated the levels of *Fmr1* mRNA from the hippocampal lysates of Wild-type and *Syngap1^+/-^* mice at PND14-16 and PND21-23. We did not observe any significant difference in *Fmr1* mRNA levels in PND14-16 (WT=0.019±0.008; *Syngap1^+/-^*=0.029±0.008; p=0.4065) and PND21-23 (WT=0.009±0.001; *Syngap1^+/-^*=0.020±0.009; p=0.3129) between Wild-type and *Syngap1^+/-^* (**Supplementary Fig 4B**). This result suggests that reduction in FMRP levels in *Syngap1^+/-^* mice at PND21-23 could be due to a decrease in *Fmr1* translation.

To further understand the translation status of *Fmr1* mRNA during PND14-16 and PND21-23, *Fmr1* mRNA present in translating fraction was quantified by qPCR. We found that *Fmr1* mRNA present in translating pool was significantly reduced in *Syngap1^+/-^* mice, as compared to Wild-type (**Fig 4C**) in PND21-23 (WT=89.34±1.03%; *Syngap1^+/-^*=73.38±4%; p=0.0257), but not in PND14-16 (WT=66.66±2.9%; *Syngap1^+/-^*=66.03±4.1%; p=0.9058) indicating reduced FMRP level as a result of decreased *Fmr1* mRNA translation in PND21-23 in *Syngap1^+/-^*.

### Altered NMDAR-mediated translation response in *Syngap1^+/-^*

Previous studies have shown increased levels of basal protein synthesis in *Syngap1^+/-^*(Wang, Held et al. 2013, Barnes, Wijetunge et al. 2015). SYNGAP1 regulates synaptic maturation during a critical time window, and our results demonstrated the altered expression of FMRP during a specific developmental stage in *Syngap1^+/-^*. Based on this, we hypothesised that the translational status could be different at these developmental stages. To study this, phosphorylation status of eukaryotic Elongation Factor 2 (eEF2) was used as a read-out of translation response. Phosphorylation of eEF2 has been shown to repress global translation (Scheetz, Nairn, & Constantine-paton, 2000). We analysed phospho/total-eEF2 in hippocampal synaptoneurosomes in response to NMDAR stimulation from Wild-type and *Syngap1^+/-^* mice at PND14-16 and PND21-23 using immunoblotting analysis. Hippocampal synaptoneurosome preparation was evaluated by validating the enrichment of PSD-95 as shown by Muddashetty *et al.*, 2007**(Supplementary Fig 5A)**. As a proof of principle, we demonstrated that NMDAR stimulation of synaptoneurosomes from Wild-type mice showed ~1.5-fold increase in phospho/total-eEF2 after 1-minute of stimulation (Basal=0.84±0.11; Stimulated=1.3±0.12; p=0.0376; **Fig 5A**).

**Figure 5.**
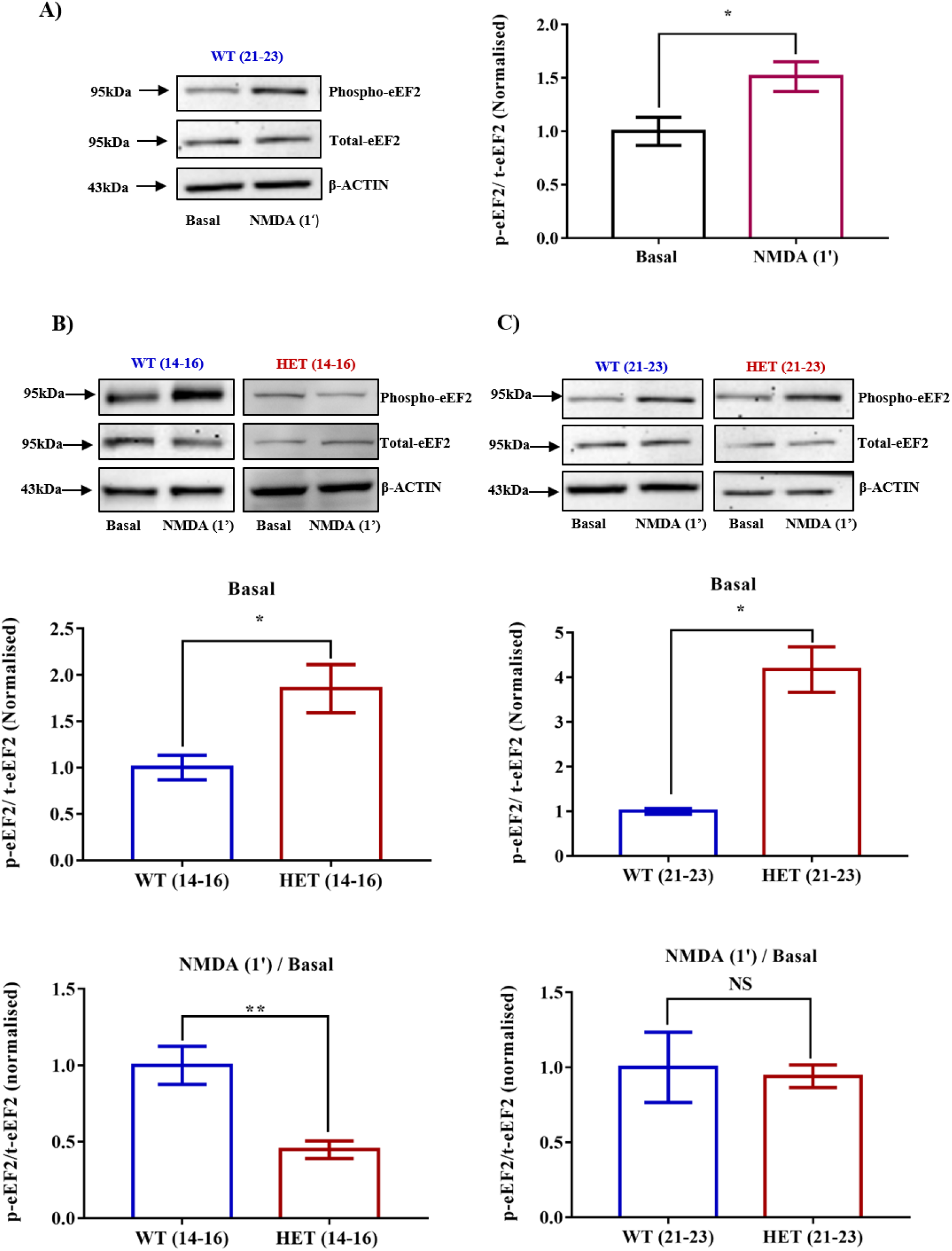
Dysregulated NMDAR mediated translation response is recovered during PND21-23 in HET. **A.** Representative immunoblot for Phospho-eEF2 and Total-eEF2 showing increased phosphorylation on NMDAR stimulation for 1-minute in synaptoneurosomes from WT during PND21-23 (*left*). Pooled data of the same represented in bar graph (*right,* Basal: n=4; Stimulated: n=4); *p<0.05. **B.** Representative immunoblots of phospho- and total-eEF2 normalised to β-ACTIN during PND14-16 in WT and HET (*top*). Bar graph shows increased phosphorylation of eEF2 at basal conditions in synaptoneurosome obtained from hippocampus of HET as compared to WT during PND14-16 in the (*middle,* WT: n=4; HET: n=3). Bar graph showing decreased phosphorylation of eEF2 in HET on NMDAR stimulation as compared to WT in PND14-16 (*below,* WT: n=4; HET: n=4). *p<0.05, **p<0.01, NS= not significant. **C.** Representative immunoblots for phospho-eEF2 and total-eEF2 normalised to β-ACTIN during PND21-23 in WT and HET (*top*). Increased phosphorylation of eEF2 at basal condition in HET synaptoneurosome as compared to WT during PND21-23 (*middle,* WT: n=3; HET: n=3). Bar graph showing the extent of phosphorylation in HET is similar to WT during PND21- 23 (*bottom,* WT: n=3; HET: n=4). *p<0.05, NS= not significant.

Further, we observed an increase in phospho/total-eEF2 in *Syngap1^+/-^* as compared to Wild-type under basal conditions in both PND14-16 (WT=0.84±0.11; *Syngap1^+/-^*=1.6±0.22%; p=0.0245) and PND21-23 (WT=0.22±0.01%; *Syngap1^+/-^*=0.9±0.11%; p=0.0233 with Welch’s correction; **Fig 5B and 5C**). We found that, at PND14-16, NMDAR-mediated increase in phosphorylation of eEF2 was lost in synaptoneurosomes in *Syngap1^+/-^* (Stimulated/Basal values for WT=1.57±0.2; *Syngap1^+/-^*=0.7±0.09; p=0.0069). Further analysis of this data by normalising to Wild-type showed a significant reduction in phospho/total eEF2 on NMDA receptor stimulation in *Syngap1^+/-^* synaptoneurosomes. (Stimulated/Basal values for WT=1.00±0.12; *Syngap1^+/-^*=0.45±0.06; **Fig 5B).** This could be because of the increased level of phosphorylated-eEF2 at basal level in *Syngap1^+/-^* mice. Surprisingly, even though we observed an increase in level of basal phospho/total-eEF2 in PND21-23 in *Syngap1^+/-^* synaptoneurosomes, NMDAR-mediated increase in phosphorylated-eEF2 in *Syngap1^+/-^* was recovered to Wild-type level (Stimulated/Basal values for WT=1.81±0.14; *Syngap1^+/-^* =1.92±0.45; p=0.8233; **Fig 5C**). A similar phenomenon was observed on 2-minute stimulation of NMDAR (**Supplementary Fig 5B, 5C**). This suggests that the NMDAR-mediated translation upon NMDAR activation at PND21-23 in *Syngap1^+/-^* returned to the level that of Wild-type.

## Discussion

Many synaptic plasticity mechanisms are dependent on activity mediated local protein synthesis in neurons (Klann, Antion et al. 2004, Pfeiffer and Huber 2006). Protein synthesis is regulated stringently in the synapse. One such crucial regulator of synaptic protein synthesis is FMRP, which is encoded by *FMR1* gene, absence of which leads to Fragile X Syndrome, a monogenic cause of ID similar to *SYNGAP1*^+/-^ (Garber, Visootsak et al. 2008, Hamdan, Gauthier et al. 2009). Our observation of enhanced mGluR-LTD in the CA1 hippocampal region of *Syngap1*^*+/*-^ is in line with the previous observation of enhanced basal protein synthesis in the heterozygous loss of *Syngap1* prompted us to investigate the role of FMRP in the pathophysiology of *Syngap1^+/-^* mutation (Wang, Held et al. 2013, Barnes, Wijetunge et al. 2015). Till date, only one report has studied interrelation between SYNGAP1 and FMRP (Barnes, Wijetunge et al. 2015). They proposed that FMRP and SYNGAP1 leads to opposite effect on synapse development, with FMRP deficits resulting in delayed synaptic maturation, and *SYNGAP1* haploinsufficiency causing accelerated maturation of dendritic spines. Considering this, Barnes et al., *Fmr1*^-/Y^ crossed with *Syngap1*^+/-^ but failed to rescue the electrophysiological deficit observed in *Syngap1*^+/-^ (Barnes, Wijetunge et al. 2015). This indicates that chronic depletion of these genes may not be an effective measure to rescue the pathophysiology observed in *Syngap1*^+/-^, as both these genes are important for normal brain development. Since SYNGAP1 is known to regulate synaptic maturation during a specific developmental window (Clement, Aceti et al. 2012, Clement, Ozkan et al. 2013) we hypothesised that the role of FMRP in *Syngap1*^+/-^ could also be developmentally regulated. Hence, we looked at the developmental expression profile of FMRP in the hippocampus of *Syngap1*^+/^ mice. Our results show reduced expression of FMRP specifically in PND21-23 in *Syngap1*^+/-^. Our study is the first to demonstrate that FMRP interacts with *Syngap1* mRNA and regulate its translation. Our result shows that the reduction in FMRP levels as well as its reduced interaction with *Syngap1* mRNA at PND21-23 in *Syngap1*^+/-^ is required for the compensatory increase in SYNGAP1 levels via increased *Syngap1* mRNA translation. In polysome profiling assay, we did not observe any significant difference in the A_254_ traces or in the distribution of protein RPLP0 between Wild-type and *Syngap1*^+/-^ animals indicating no difference in the basal translation in the hippocampus from *Syngap1*^+/-^ animals at PND14-16 and PND21-23.

Studies have reported that NMDAR-mediated signalling is dysregulated in *Syngap1*^+/-^ (Komiyama, Watabe et al. 2002, Rumbaugh, Adams et al. 2006, Carlisle, Manzerra et al. 2008). These studies have further shown that SYNGAP1 associates with NR2B (Rockliffe and Gawler 2006) and negatively regulates NMDA receptor-mediated ERK activation (Kim, Dunah et al. 2005) and, hence, regulates insertion of AMPAR in the post-synaptic membrane (Rumbaugh, Adams et al. 2006). In line with this, studies have demonstrated increased basal levels of ERK phosphorylation in *Syngap1*^+/-^ (Komiyama, Watabe et al. 2002) which does not explain the deficits observed in NMDAR-LTP in *Syngap1*^+/-^ mice as NMDAR stimulation resulted in a robust increase in ERK activation in slices from *Syngap1*^+/-^ mice (Komiyama, Watabe et al. 2002). Thus, to understand the deficits seen in NMDAR-mediated signalling in *Syngap1*^+/-^ mice, we studied NMDAR-mediated translation repression. It has already been reported that NMDAR activation causes a reduction in global translation through phosphorylation of eEF2 (Scheetz, Nairn et al. 2000). In our study, we measured the basal levels of phosphorylated eEF2 in hippocampal synaptoneurosomes from Wild-type and *Syngap1*^+/-^ at PND14-16 and PND21- 23 which showed increased phosphorylation of eEF2 at the basal condition in *Syngap1*^+/-^ in these age groups. This increase in the basal level of phosphorylation of eEF2 could be due to enhanced excitatory neuronal activity in *Syngap1*^+/-^ which might lead to an increase in Ca^2+^ levels and a subsequent increase in eEF2 phosphorylation via Ca^2+^-Calmodulin kinase. We report that, at PND14-16, NMDAR activation fails to cause eEF2 phosphorylation in *Syngap1*^+/-^ animals. Strikingly, at 3-week of age, even though we observed an increase in basal phospho/total-eEF2 in *Syngap1^+/-^* synaptoneurosomes, NMDAR-mediated increase in eEF2 phosphorylation was similar to Wild-type (p=0.8233; **Figure 5C**). This observation suggests that NMDAR-mediated translation response at PND21-23 in *Syngap1^+/-^* is restored. This change observed in 3-week old mice could be due to a compensatory mechanism through increased NMDAR signalling. These findings further corroborate with the observations made by Clement et al. in which they have demonstrated increased synaptic transmission and increased AMPA/NMDA in PND14, but returns to normal level in the later age (Clement, Aceti et al. 2012). Based on our findings, we propose a model in which increased NMDAR-mediated protein synthesis is compensating for the loss of SYNGAP1 during development in *Syngap1^+/-^* (**Fig 6**). We further propose that fine-tuned downregulation of *Fmr1* translation during a specific developmental window in *Syngap1*^+/-^ mice might compensate for the dysregulation in NMDAR-mediated signalling.

**Figure 6:**
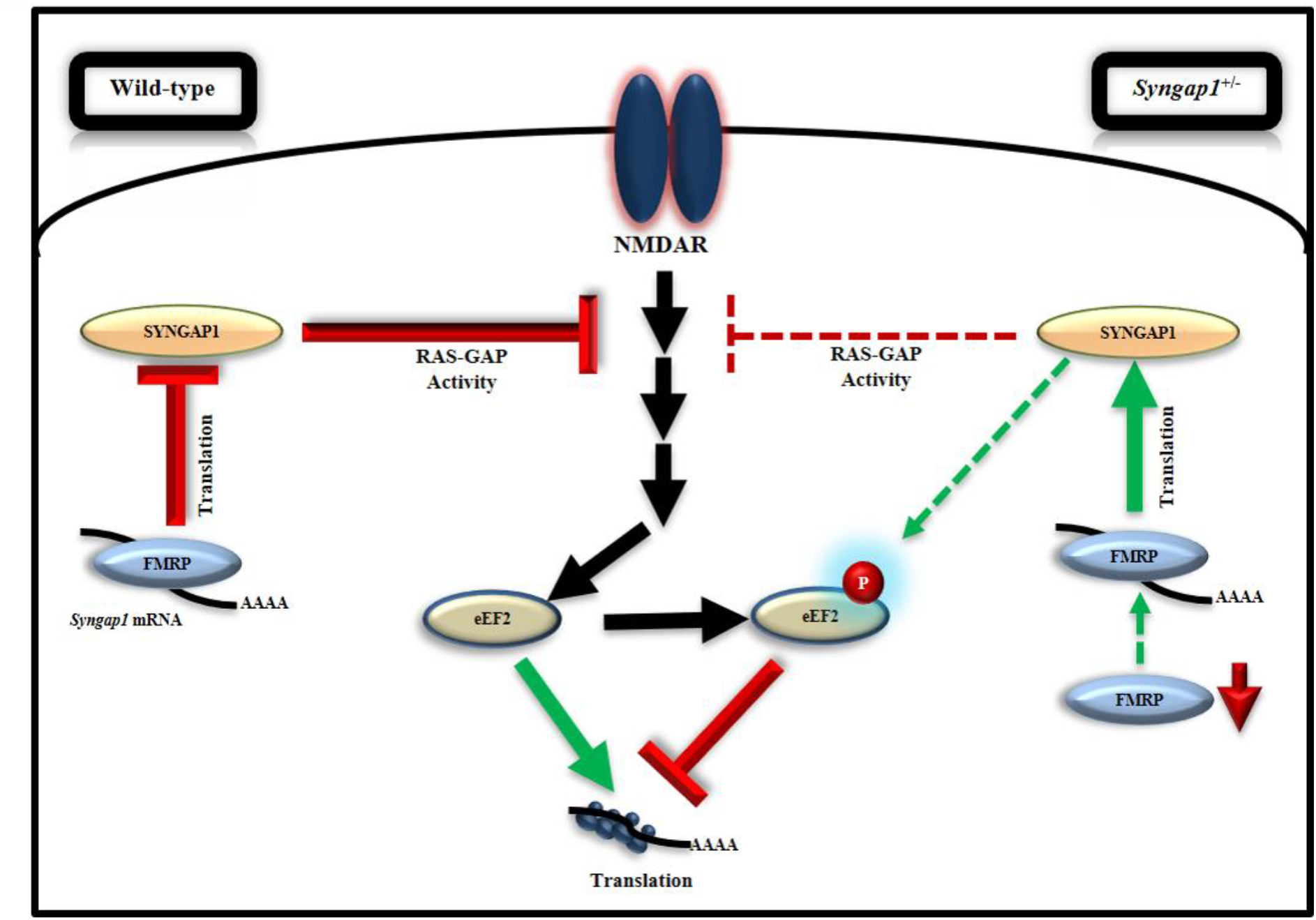
Model illustrating the regulation of FMRP-mediated translation of *Syngap1* during development: This model shows that FMRP regulates *Syngap1* mRNA translation, which in turn regulates NMDAR signalling. In WT, NMDAR stimulation in synapse leads to increased phosphorylation of eEF2, which results in global translation inhibition and the signalling is efficiently regulated by SYNGAP1. Whereas, in *Syngap1^+/-^* at PND14-16, NMDAR mediated signalling is impaired depicted by the loss of phosphorylation response to eEF2, as the level of SYNGAP1 is low. At PND21-23 in *Syngap1^+/-^*, FMRP level is low which allows increased translation of *Syngap1* mRNA leading to an increased SYNGAP1 level compared to PND14-16. Thus, an elevated level of SYNGAP1 might recover the NMDAR mediated signalling via phosphorylation of eEF2.

In conclusion, our study suggests that altered response to activity mediated protein synthesis during development is one of the major causes of abnormal neuronal function in *Syngap1^+/-^*. However, chronic depletion of two genes with common core pathophysiology may not be an effective measure to rescue the deficits observed in either of these mutations *i.e*. *Fmr1^-/y^* and *Syngap1^+/-^*, as both these genes are important for normal brain development. Therefore, modulating these proteins at a certain developmental window could be a potential therapeutic strategy for treating ID-related pathophysiology.

## Acknowledgement

This work was supported by grants to JPC by DST-SERB (SB/YS/LS-215/2013), and to RSM in part, by Dept. of Biotechnology, India (BT/PR8723/AGR/36/776/2013, and BT/IN/Denmark/07/RSM/2015-2016), and intramural funds from both the Institutes. We Sincerely thank Prof Gavin Rumbaugh for providing us *GFP-Syngap1* construct. We convey our gratitude to Dr. Hashim Reza for technical support in Figure 2. We thank Bhavana Kayyar, Utsa Bhaduri, and Vijay Kumar M J for technical support in our bioinformatics analysis, and Sudhriti Ghosh Dastidar for technical advice on Figure 4. Further, we thank Dr Ravi Manjithaya and his group, and Prof. Kaustuv Sanyal’s group for technical assistance. We also thank Prof MRS Rao, Prof Tapas K Kundu, Dr Sheeba Vasu, Dr Sourav Banerjee, Dhriti Nagar, Natasha Basu, and JPC and RSM lab members for critical comments on this manuscript. We thank Dr Prakash, Animal house in-charge, JNCASR, for help with the animal facility.

## Author Contribution

AP and BN performed all the experiments. JPC did mGluR-LTD in Figure 1A. SS performed part of the experiments in Figure 1B and 1D. AP, BN, RSM, and JPC designed the experiments and wrote the manuscript. SS edited the manuscript.

---

**Supplementary Figure 1:**
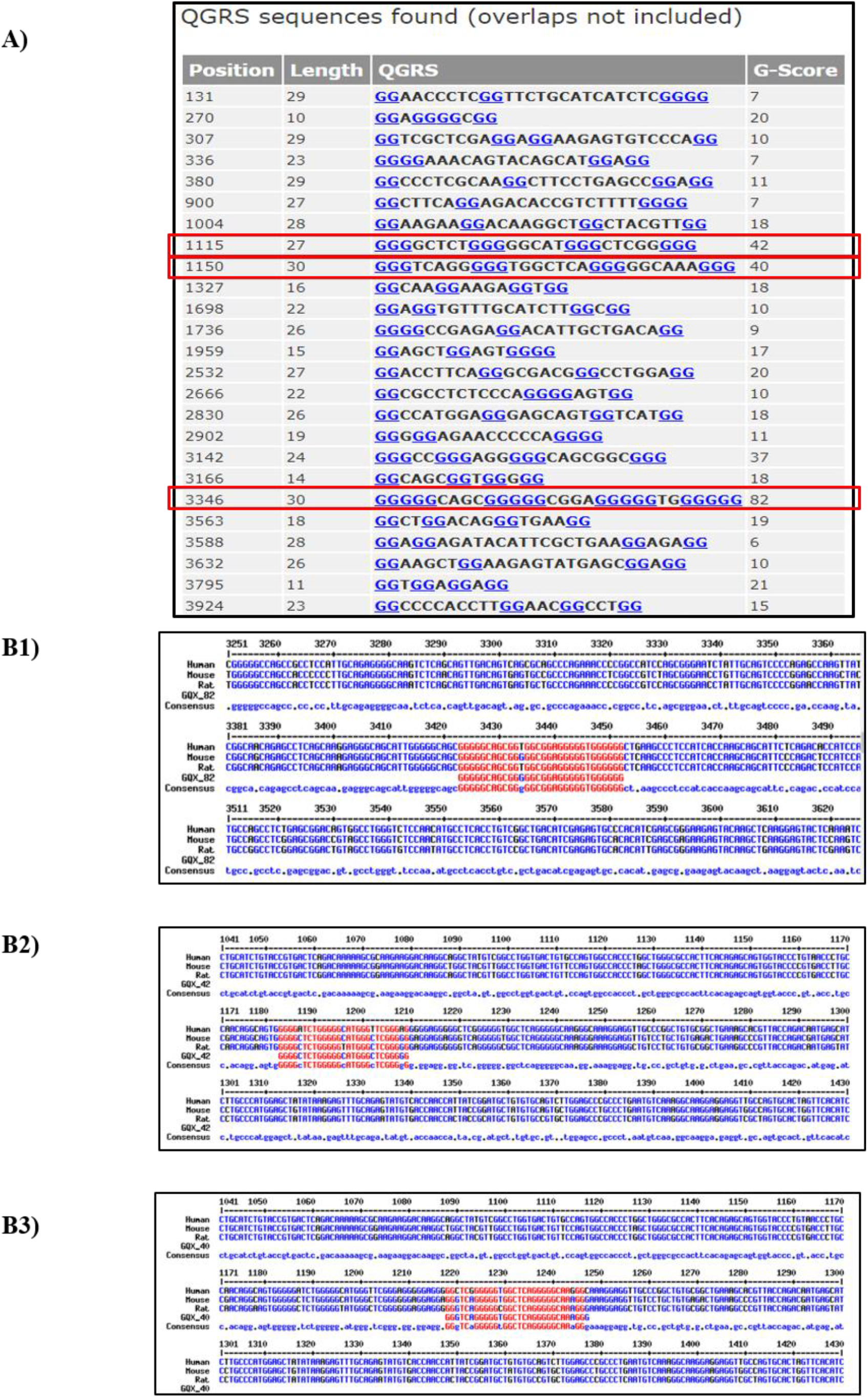
*Syngap1* mRNA contains predicted G quadruplexes. **A.** Multiple putative G-quadruplex was detected using QGRS Mapper in the validated sequence available for mouse *Syngap1* from NCBI (Gene ID: 240057). Three high G-score having G-quadruplex sequences are highlighted in the red box. All these sequences have been mapped in the Coding Sequence (CDS). **B1-3.** Multiple sequence alignment of the three high score G-quadruplexes of mouse *Syngap1* compared with Human and Rat. G scores: 82 (B1), 42 (B2), and 40 (B3) showing putative G-quadruplexes conserved among Human, Mouse, and Rat respectively.

**Supplementary Figure 2:**
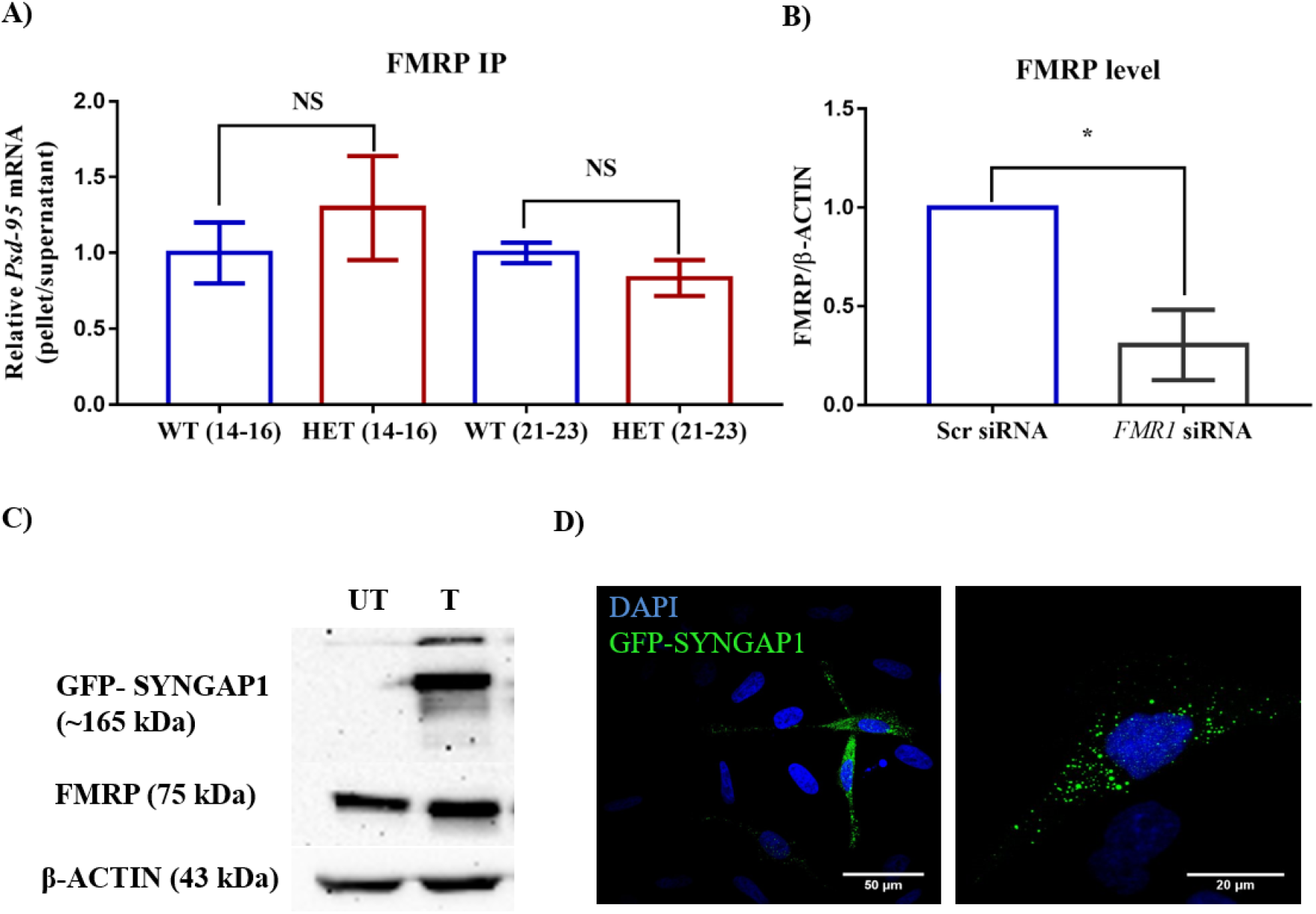
FMRP interacts with *Syngap1* mRNA in the hippocampus. **A.** Bar graph showing relative *Psd-95* mRNA enrichment in FMRP IP pellet compared to supernatant from hippocampus at PND14-16 (WT: N=7; HET: N=3) and PND21-23 (WT: N=5; HET: N=4) normalised to WT. Unpaired Student’s *t*-test. NS=not significant. **B.**Bar graph shows reduced level of FMRP in the *FMR1* siRNA treated cells compared to *scr* siRNA treated control (WT: N=4; HET: N=4). Unpaired Student’s *t*-test; * p<0.05. **C.** Representative immunoblot for SYNGAP1 and FMRP showing the expression of SYNGAP1 in transfected (T) compared to Un-transfected (UT) control **D.** Representative images of Hela cells showing the expression of GFP-SYNGAP1 (Green). Cell nucleus are stained with DAPI (Blue). Right panel shows higher magnification image where GFP-SYNGAP1 shows punctate structure.

**Supplementary Figure 3:**
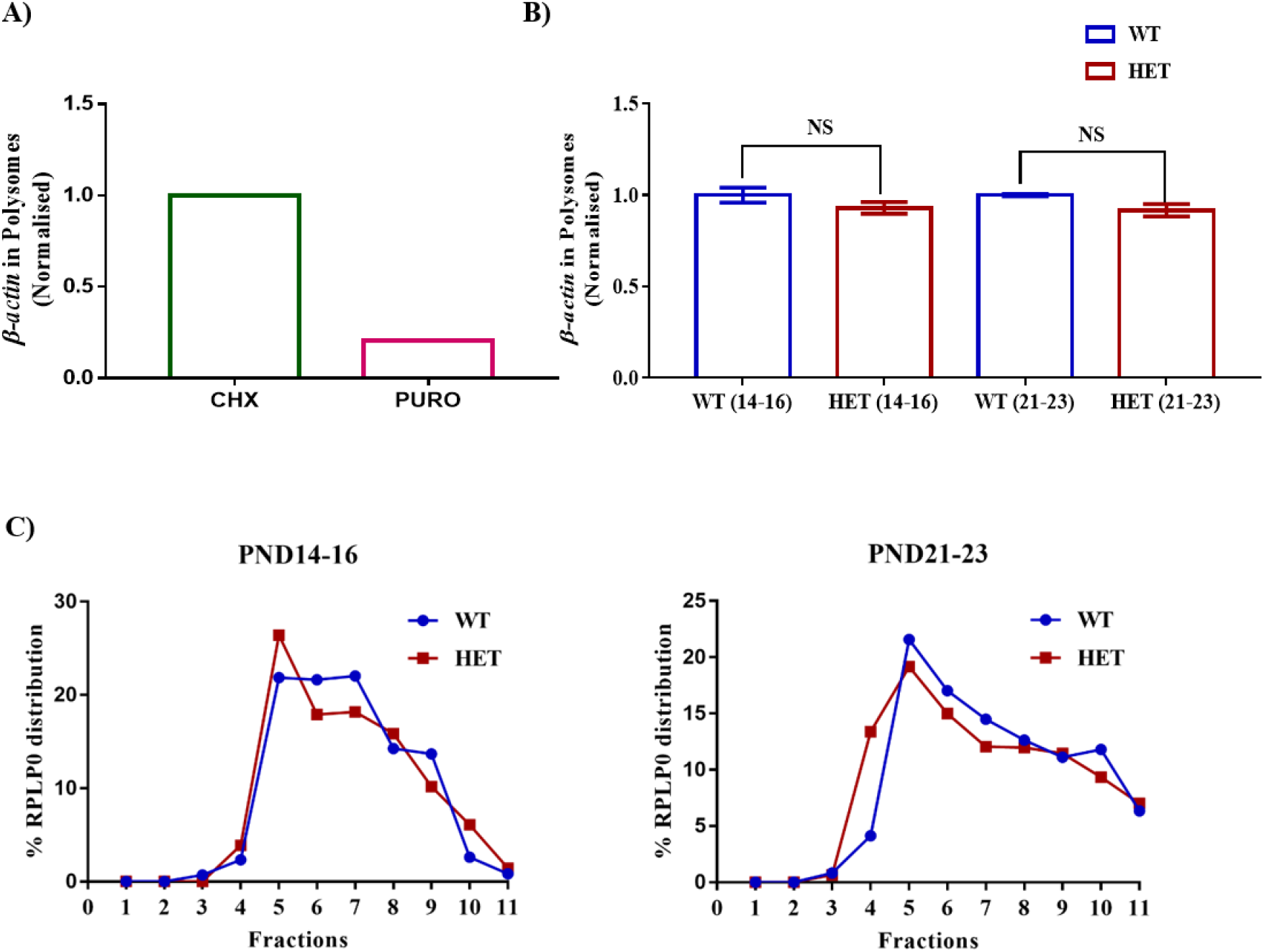
RPLP0 distribution unaltered in polysome. **A.** *β-actin* mRNA distribution in polysome treated with cycloheximide and puromycin **B.** Bar diagram showing *β-actin* mRNA distribution in Cycloheximide treated polysome HET normalised to WT in PND14-16 (WT: n=6; HET: n=6) and PND21-23 (WT: n=4; HET: n=5). NS = not significant. **C.** Representative percentage RPLP0 distribution line graph in PND14-16 (*left*) and PND21- 23 (*right*).

**Supplementary Figure 4:**
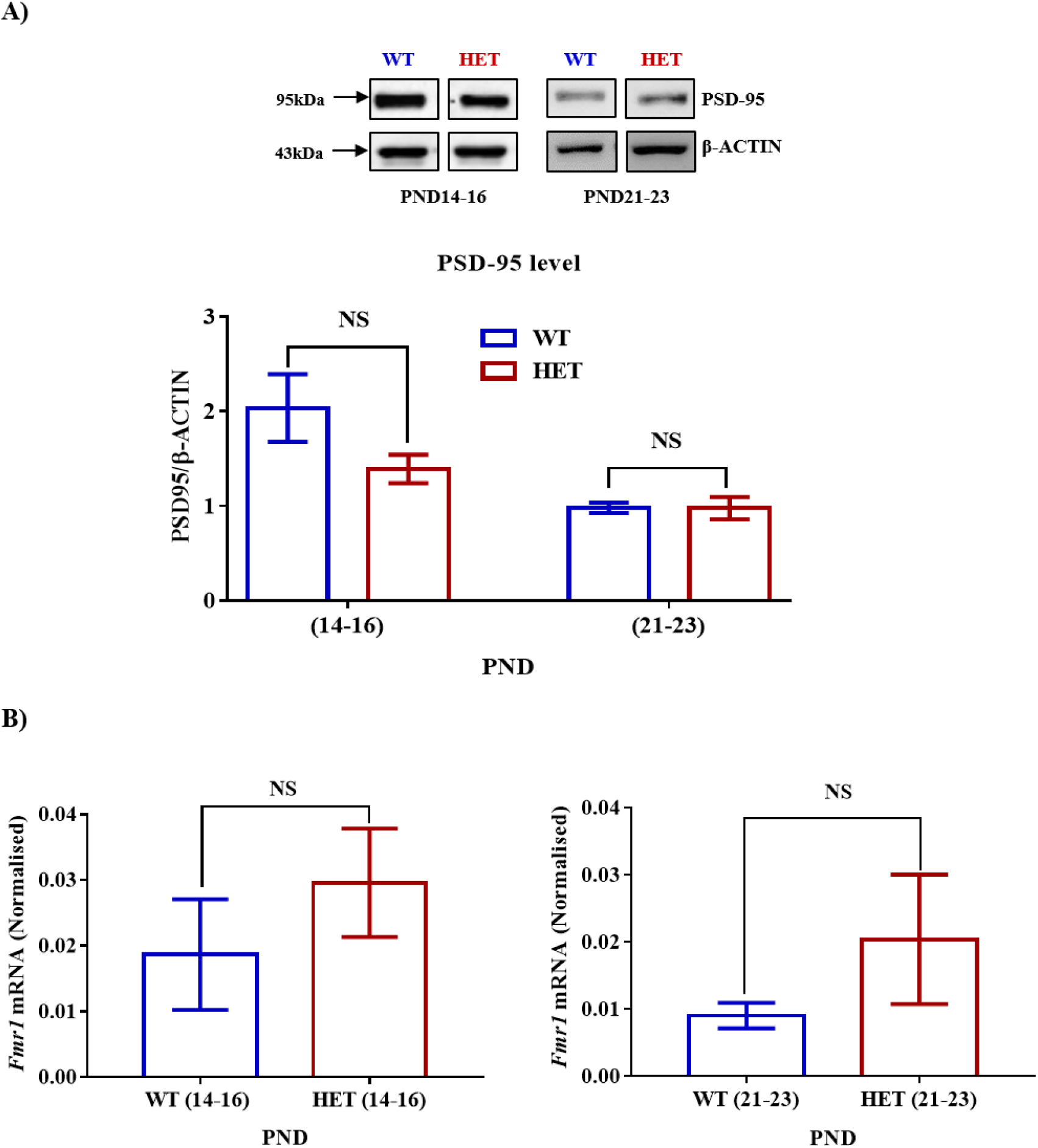
PSD-95 level in PND14-16 and PND21-23. **A.** Representative immunoblots for PSD-95 normalised to β-ACTIN in the hippocampus during PND14-16 and PND21-23 in WT and HET. *Below* Bar graph showing a no significant difference in the level of PSD-95 at PND14-16 (WT: n=6; HET: n=6) and PND21-23 (WT: n=10; HET: n=4) between WT and HET. *p<0.05, NS= not significant. **B.** Bar graph depicting relative *Fmr1* mRNA normalised to *β-actin* from total hippocampal lysate at PND14-16 (*left,* WT: n=3; HET: n=3) and PND21-23 (*right,* WT: n=3; HET: n=3); NS= not significant.

**Supplementary Figure 5:**
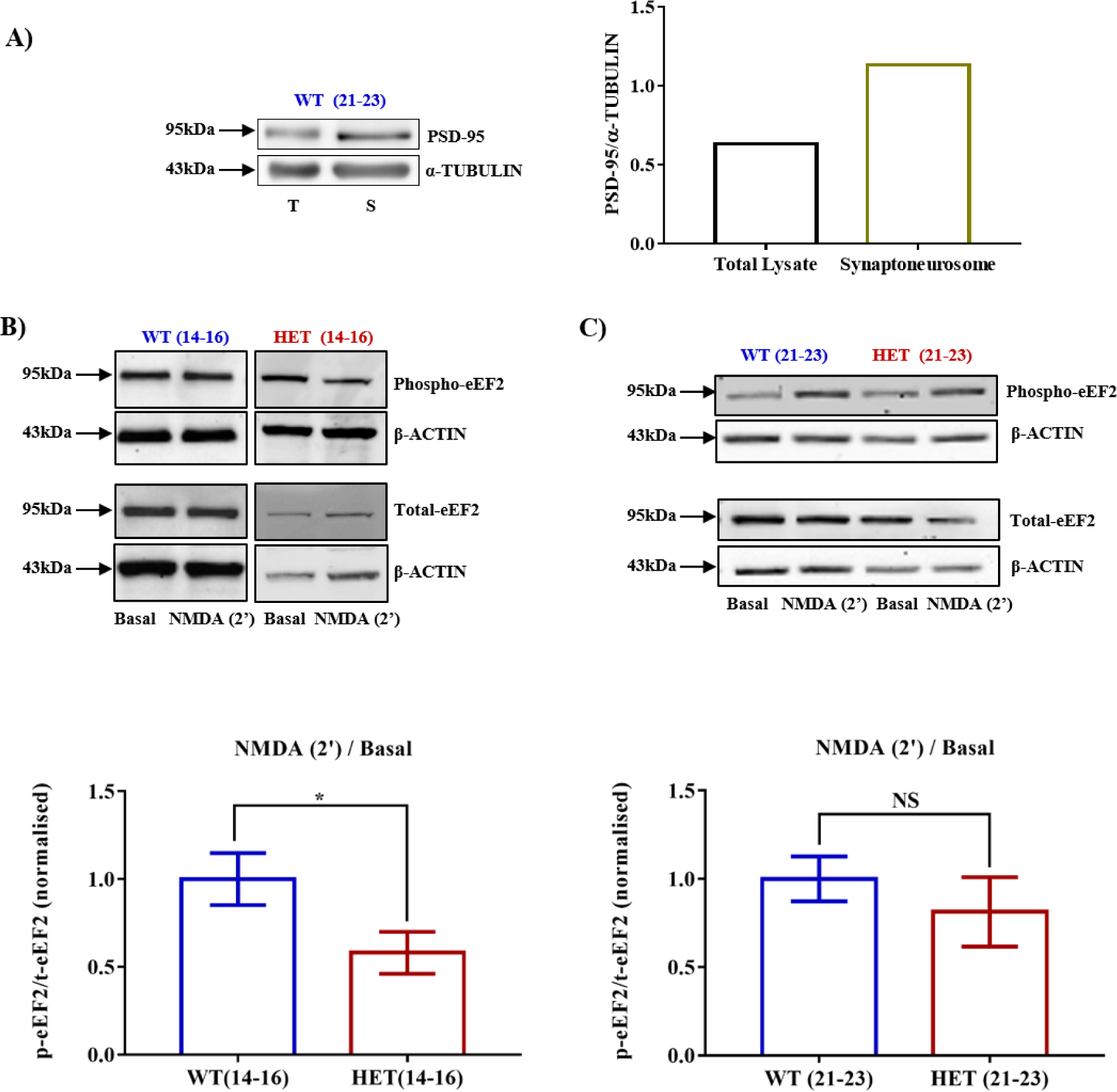
Altered phosphorylation of eEF2 in HET. **A.** Representative immunoblot depicting the enrichment of PSD-95 in synaptoneurosome (S) compared to Total hippocampal lysate (T; *left*). Bar graph showing quantified data normalised to α-TUBULIN (*right*). **B.** Representative immunoblot images for Phospho-eEF2, Total-eEF2, and β-ACTIN synaptoneurosomes after 2-minute NMDAR stimulation during PND14-16 (*top*). Bar graph showing that 2-minute activation of NMDAR alters phosphorylation of eEF2 in HET (n=3) compared to WT (n=3) during PND 14-16 (*below*). *p<0.05, NS= not significant **C.** Representative immunoblot for Phospho- and Total-eEF2 normalised to β-ACTIN synaptoneurosomes stimulated with NMDA for 2-minute in WT and HET during PND21-23 (*top*). Bar graph depicting unaltered phosphorylation of eEF2 in HET (n=3) compared to WT (n=3) post-2-minute activation of NMDAR. NS= not significant.

